# POSH (Plenty of SH3): a novel genetic modifier of Myc function in Drosophila

**DOI:** 10.64898/2026.10.19.760028

**Authors:** Vicky Kuo, Margaret Daniel, Sheri Zola, Paola Bellosta

## Abstract

MYC is a conserved transcriptional regulator that coordinates cellular growth, metabolism, and tissue homeostasis through a broad network of downstream effectors. However, the signaling components that translate MYC activity into specific cellular responses remain incompletely defined. Here, using *Drosophila melanogaster*, we identify Plenty of SH3s (POSH), a conserved RING-domain scaffold protein and regulator of JNK signaling, as a novel genetic interactor of Myc. POSH was identified in a genetic screen for modifiers of a Myc-dependent phenotype in the developing eye. The interaction was independently confirmed using a *UAS-Posh* transgene. Quantitative analysis showed that reduced Myc primarily decreased ommatidial size, whereas POSH expression predominantly reduced ommatidial number, with this effect becoming more pronounced in Myc-sensitized backgrounds. Myc overexpression also increased *Posh* transcript levels, and the *Posh* regulatory region contains a canonical E-box motif, supporting a functional relationship between the two genes while leaving open the question of whether *Posh* is a direct Myc transcriptional target. Co-expression experiments further revealed a strong morphological interaction between Myc and POSH in the adult eye. Together, these findings identify POSH as a previously unrecognized genetic modifier of Myc function in *Drosophila* and provide a framework for investigating how Myc interfaces with stress-responsive signaling pathways during tissue development.

## 1. Introduction

MYC is an evolutionarily conserved transcription factor that coordinates cellular growth, proliferation, metabolism, protein synthesis, and stress responses. In *Drosophila*, Myc is encoded by the single *myc/diminutive* (*dm*) gene and retains the major biochemical and biological properties of its vertebrate counterparts [1,2]. Similar to mammalian MYC, *Drosophila* Myc heterodimerizes with Max and binds E-box sequences to regulate a broad spectrum of target genes, including genes involved in ribosome biogenesis, protein synthesis, metabolism, and cellular stress responses [3][4]. Reduced Myc activity impairs cellular and organismal growth, resulting in small flies (*diminutive*) [5], whereas increased Myc expression promotes growth, promoting ribosomal biogenesis [3,4,6-8]. Myc also integrates nutrient-dependent signals with growth control downstream of insulin/TOR signaling, including non-cell-autonomous regulation of systemic growth via its expression in the larval fat body [9-11].

The biological effects of Myc depend on cellular context and relative Myc activity, and extend beyond growth. Elevated Myc levels can induce apoptosis [12,13] or autophagy [14,15]. In developing tissues, differences in relative Myc activity can trigger cell competition, in which cells with lower Myc activity are eliminated by better-fitted neighboring cells [8,16]. These observations provided the rationale for generating flies with a Myc-sensitized background, allowing us to identify *Plenty of SH3s (Posh*) in a genetic screen as a modifier of the Myc-dependent growth phenotype *in vivo*.

POSH is a conserved RING-domain scaffold protein involved in several stress signaling pathways. In *Drosophila*, POSH has been shown to participate in JNK signaling and stress-induced, caspase-dependent apoptosis [17-20]. POSH has also been reported to regulate the Hippo pathway by promoting the ubiquitination and degradation of the target Expanded (Ex), thereby modulating Yorkie-dependent tissue growth and regeneration [21].

These findings link POSH to pathways controlling apoptosis and tissue growth, providing a rationale for investigating its genetic interaction with Myc.

## 2. Materials and Methods

### Fly Husbandry and lines

Flies were raised at low density, at 25 ^°^C on a standard food medium containing 6 g/L agar, 75 g/L corn flour, 100 g/L white sugar, 50 g/L fresh yeast, and 10 g/L inactivated yeast powder, along with nipagin and propionic acid (Acros Organic). The following *Drosophila* stocks were obtained from the Bloomington Drosophila Stock Center (BDSC): *tubulin-GAL4* (B5138), B17054, B17081, B17108, and B16997 (see Table 1). *GMR-GAL4*, a kind gift from Peter Gallant (Julius-Maximilians-Universität Würzburg, Germany), *UAS-HA-Myc* [22], *w,myc*^*P0*^ *tub-FRT>(dmyc-cDNA)>FRT GAL4 ey-FLP/Y* and control *w, tub-FRT>(dmyc-cDNA)>FRT GAL4 ey-FLP/Y* [22], *UAS-Posh* [20].

**Table 1.** Candidate genes identified in the genetic modifier screen. EP lines that reproducibly modified the Myc-dependent eye phenotype are listed with their BDRC stock numbers, chromosomal locations, associated genes, full gene names, and reported biological functions.

| EP line | BDRC | Gene/Ch | Full gene name | Reported function |
| --- | --- | --- | --- | --- |
| *EP-2418* | 17054 | *NimC3/2L* | *Nimrod C3(hll)* | Nimrod-family cell-surface protein involved in Innate immunity and phagocytosis |
| *EP-2582* | 17071 | *robo2 (lea)/2L* | *roundabout 2 / leak* | Transmembrane receptor involved in axon guidance and developmental signaling |
| *EP-3291* | 17108 | *Arts/3R* | *Apoptosis-related TGF-β signaling* | Regulation of apoptosis and cell-death signaling |
| *EP-1206* | 16997 | *POSH/2R* | *plenty of SH3s* | SH3-domain/RING-containing scaffold protein associated with JNK and apoptotic signaling |

### Genetic Modifier Screen

A collection of 162 P(EP) insertion lines from the Rørth EP controlled-misexpression collection was obtained from the Bloomington Drosophila Stock Center (BDSC, Indiana University, USA) and used in a dominant genetic modifier screen. P(EP) elements contain GAL4-responsive UAS sequences that can drive expression of genes adjacent to the insertion site, depending on the insertion site’s position and orientation. Males F1 progeny carrying one EP insertion were screened for dominant modification of the adult eye phenotype in the Myc-sensitized background (to females of the previously described *ey-FLP*-based Myc tester stocks (*w,myc*^*P0*^ *tub-FRT>(dmyc-cDNA)>FRT GAL4 ey-FLP/FM7* and control *w, tub-FRT>(dmyc-cDNA)>FRT GAL4 ey-FLP/FM7*) [22]. Males F_1_ progeny carrying one EP insertion were screened for dominant modification of the adult eye phenotype in males of the Myc-sensitized background (*ey>dmyc*^*P0*^*/Y; EP-line/+*) relative to the corresponding control (*ey>dmyc*^*WT*^*/Y; EP-line/+*). Candidate modifiers were retested in independent crosses. The EP-1206 insertion (B16997), expressing *plenty of SH3s (POSH*), was selected for further characterization.

### Adult Eye and Ommatidial Analysis

Three-day-old adult male flies were analyzed using two genetic systems. For experiments involving reduced Myc activity, the *ey-FLP*-based *dmyc* lines previously described [22]. In these lines, FLP-mediated recombination under the *eyeless* compartment during development removes *dmyc* cDNA and simultaneously permits GAL4 expression. For Myc and POSH overexpression experiments, *GMR-GAL4* was used to drive UAS transgenes in the developing eye.

Adult flies were collected in Eppendorf tubes containing 1 ml of 80% acetone/20% water (v/v) and stored at -80 °C before fixation and preparation for scanning electron microscopy (SEM), as previously described [22]. SEM images of one eye per animal, a minimum of 5 animals per genotype, were used for quantitative analysis. Total ommatidial number was determined by counting all ommatidia in each eye, in the image, whereas ommatidial size was determined from the same image by measuring the area of 20 ommatidia located in the central region of the eye using Adobe Photoshop CS4 (Adobe Systems Inc., San Jose, CA, USA), following the previously established procedure. The measurements from the 20 ommatidia were averaged to obtain a single value for each animal, with each animal considered an independent biological replicate. The number of animals analyzed for each genotype is indicated in the corresponding graphs in Figures 2 and 4. Data are presented as mean ± SD. Statistical significance was assessed by one-way ANOVA followed by Sidak’s multiple-comparisons test for the indicated pairwise comparisons, with *P* < 0.05 considered statistically significant.

### RNA Extraction and Quantitative RT-PCR

RNA was extracted from whole larvae ubiquitously expressing the EP-1206 insertion under the control of *tubulin-Gal4*, with *tubulin-Gal4* alone as the control (Figure 1F); or from eye imaginal discs expressing *UAS-Myc* using the *GMR-Gal4*, with *GMR-Gal4* alone as a control (Figure 4A). Total RNA was isolated using the Qiagen-RNeasy mini kit; cDNA was synthesized from 1µg of total RNA using Superscript IV-VILO (#11754050 Invitrogen). Quantitative PCR was performed using PowerTrack SYBR Green Master Mix (Thermo Fisher Scientific). Transcript abundance was normalized to rp49, and expression levels are reported as fold change relative to the corresponding control. Experiments were repeated twice, with an average of six whole larvae and about 30-40 eye-imaginal discs per biological replicate per genotype.

**Figure 1.**
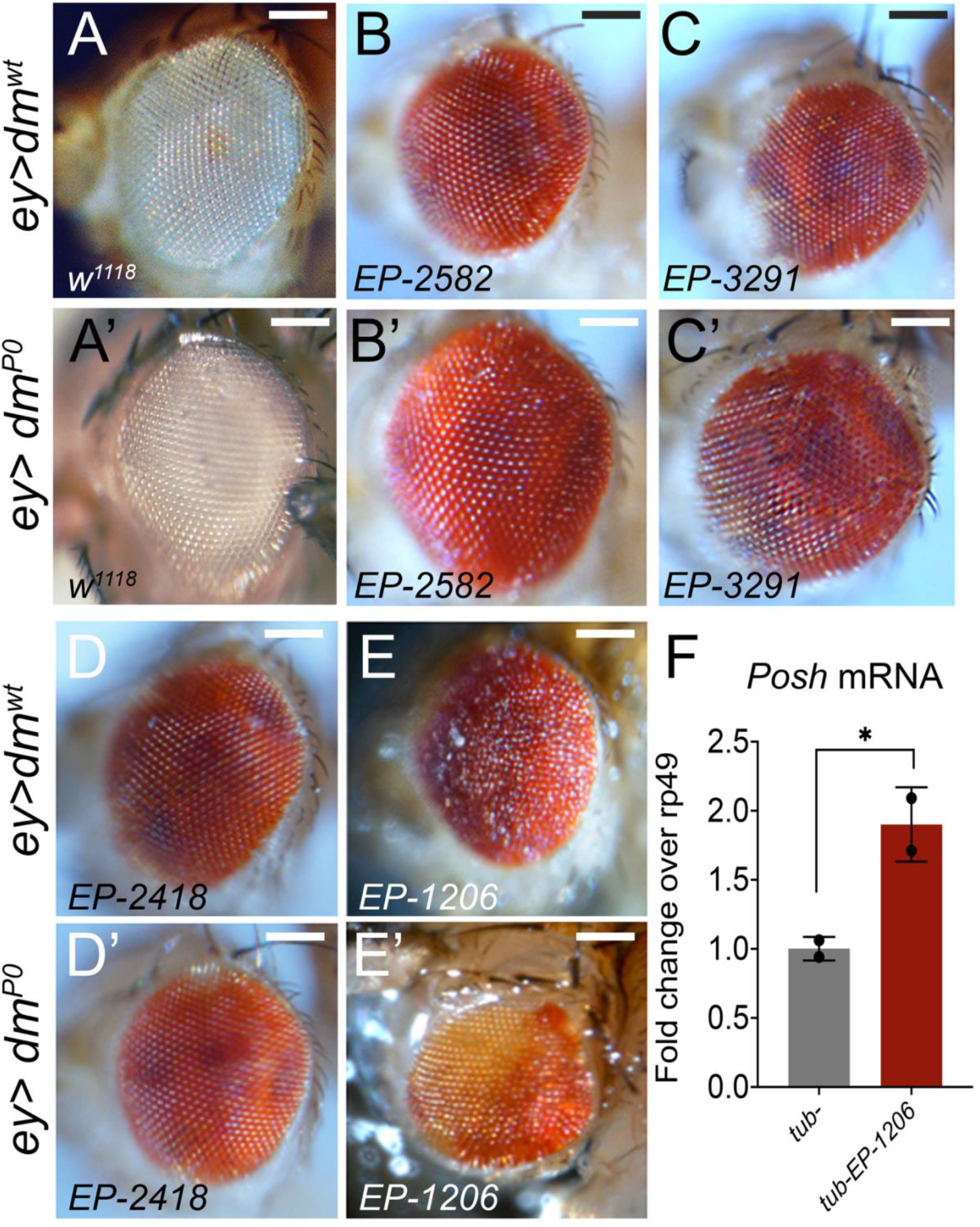
Identification of genetic modifiers of the Myc-dependent adult eye phenotypes. (A-E’) Representative adult eyes from control flies ey>*dm*^*WT*^ (A-E) and Myc hypomorphic mutant background ey>*dm*^P0^ (A′-E′). (A,A’) *w*^*1118*^ control, (B,B′) EP-2582, (C,C′) EP-3291, (D,D’) EP-2418, and (E,E) EP-1206. Scale bars, 100 µm. (F) RT-qPCR analysis of *Posh* mRNA in larvae expressing EP-1206 under the *tubulin* promoter, compared to *tubulin w*^*1118*^ control. *POSH* transcript levels were normalized to *rp49* and expressed as fold change relative to the control. Bars represent mean ± SD; dots indicate independent biological experiments ^*^*p* < 0.05.

Primer sequences were as follows: POSH forward-GCAGACGCCCAGATGGCACAA; reverse-GCCTGCTGCTGACGAGGTGG, rp49 forward-GACGCTTCAAGGGACAGTATCTG; reverse-AAACGCGGTTCTGCATGAG.

## 3. Results

### 3.1 Screen for Dominant Genetic Modifiers of a Myc-Dependent Phenotype

To identify genes that genetically interact with *myc in vivo*, we performed a dominant modifier screen using 162 P(EP) insertion lines from the Rørth EP controlled-misexpression collection [23,24]. P(EP) elements contain UAS sequences that can drive expression of genes adjacent to the insertion site in the presence of GAL4, allowing identification of genes whose altered expression modifies a sensitized phenotype. The screen used the *ey-FLP*-based tester line *ey>dm*^*P0*^ and *ey>dm*^*wt*^, which provide an in vivo readout of Myc-dependent eye growth [22]. In this system, FLP-mediated recombination under the control of the *eyeless* compartment removes a rescuing *dmyc-cDNA* expressed from the *tubulin* promoter while simultaneously permitting GAL4 expression. Individual EP insertions were therefore tested in control *dmyc-WT* or *dmyc*^*P0*^-sensitized tissue to determine whether they modify the adult eye phenotype associated with reduced endogenous Myc function. F1 progeny carrying the Myc tester system and one copy of each EP insertion were scored for dominant enhancement or suppression of the adult eye phenotype.

Of the 162 EP lines screened, four reproducibly modified the Myc-dependent eye phenotype, representing approximately 2.5% of the collection (Table 1 and Figure 1). The phenotypes associated with EP-2418 (Figure 1B,B′), EP-2582 (Figure 1C,C′), EP-3291 (Figure 1D,D′), and EP-1206 (Figure 1E,E′) were compared with the corresponding controls (Figure 1A,A′). EP-1206, associated with the *Posh* gene (plenty of SH3s), was selected for further characterization because of its pronounced modifier phenotype. In addition, the 5’ promoter region of *Posh* contains a canonical E-box motif that could function as a potential Myc-binding sequence [3], although its functional significance remains to be established. Quantitative RT-PCR from whole larvae showed that *tub>EP-1206*-driven expression increased *Posh* transcript levels approximately 1.9-fold relative to *control tub*-*w*^*1118*^ (Figure 1F).

### 3.2 Expression of POSH Modifies the Eye Phenotype in Myc-Sensitized Background

*Drosophila* POSH is an evolutionarily conserved scaffold protein involved in cellular stress responses, including apoptosis [19]. It is an 838-amino-acid protein characterized by an N-terminal RING finger domain, associated with E3 ubiquitin ligase activity, and four SH3 domains that mediate protein-protein interactions (Figure 2). Its structure shares a similar domain organization with human homologs SH3RF1 (POSH1) and SH3RF3 (POSH2) [25]. Although the overall amino acid sequence identity between *Drosophila* and mouse POSH is relatively low (approximately 24%), the functional domains are more highly conserved (43-67% sequence similarity), including the activation of JNK signaling, suggesting conservation of their molecular functions across species [17,26].

**Figure 2.**
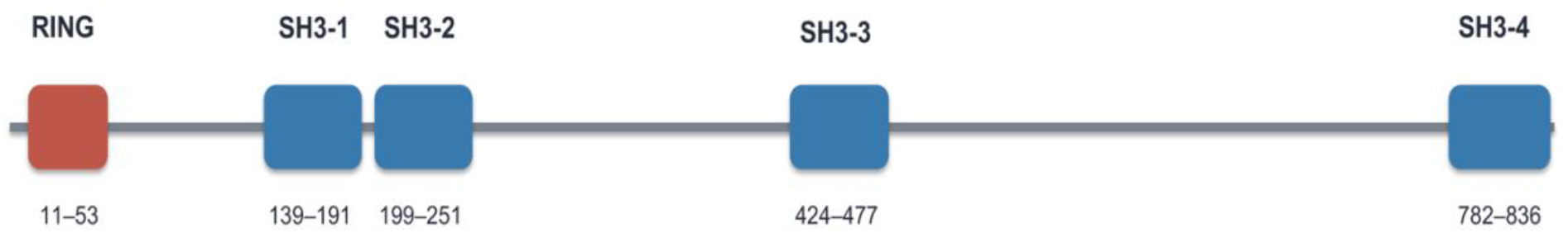
Domain organization and evolutionary conservation of *Drosophila* POSH. POSH contains an N-terminal RING finger domain and four SH3 domains involved in protein-protein interactions.

To validate the genetic interaction identified in the EP screen and test whether it could be reproduced independently of the insertion, we expressed POSH from a *UAS-Posh* transgene in a background with different levels of Myc activity. In the *dm*^*wt*^ background, POSH overexpression produced a modest alteration in adult eye morphology (Figure 3A-B). Total ommatidial number decreased from 726 ± 17 to 621 ± 45, with a smaller change in mean ommatidial area (239 ± 11 versus 225 ± 10 μm^2^). In contrast, reduced Myc activity markedly enhanced the POSH-associated phenotype. *UAS-Posh* expression in the hypomorphic *dm*^*P0*^ background produced smaller, disorganized eyes (Figure 3A’-B’), and the phenotype was more pronounced in the in *dm*^*4*^ null background (Figure 3A”-B”). This enhancement was mainly associated with reduced ommatidial number, from 735 ± 16 to 560 ± 60 in the *dm*^*P0*^ flies and from 650 ± 44 to 442 ± 116 in the *dm*^*4*^ mutant flies. By comparison, ommatidia are changed little in the *dm*^*P0*^ background (217 ± 9 versus 215 ± 20 μm^2^), (Figure 3B-C).

**Figure 3.**
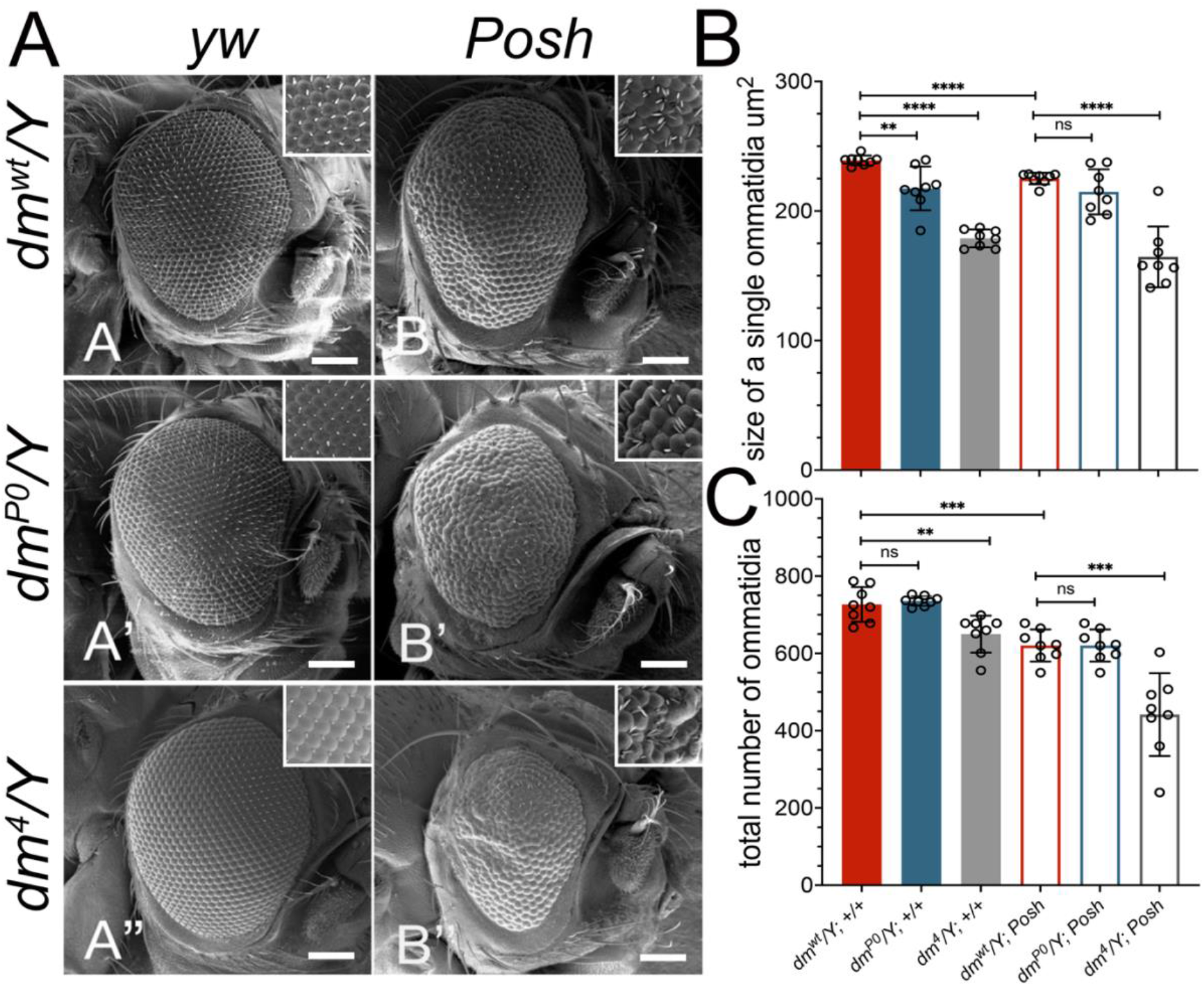
POSH modifies adult eye morphology in Myc-sensitized backgrounds. Representative scanning electron micrographs of control eyes from *dm*^*wt*^*/Y* (A), hypomorphic allele *dm*^*P0*^ (A′), and the null-mutant *dm*^*4*^ (A″) background, and the corresponding eyes expressing *UAS-Posh* (B, B’, B”). Insets show higher-magnification views of the ommatidial organization. (C) Quantification of mean ommatidial area and total ommatidial number is shown on the right. Reduced Myc activity was associated primarily with smaller ommatidia, whereas POSH overexpression had a comparatively modest additional effect on ommatidial size. In contrast, POSH overexpression reduced ommatidial number more strongly as Myc activity decreased. Data are mean ± SD; individual points represent independent eyes. Statistical significance is indicated in the graphs; ns, not significant; ^**^*p* < 0.01, ^***^*p* < 0.001, ^****^*p* < 0.0001. Scale bar, 100 μm.

Thus, reduced Myc activity primarily affected ommatidial size, whereas POSH overexpression in Myc-sensitized background predominantly reduced ommatidial number. These experiments independently reproduced the interaction observed with EP-1206 and identify POSH as a genetic modifier of the Myc-dependent eye phenotype.

### 3.3 Myc Overexpression Increases *Posh* Transcript Levels and Genetically Interacts with POSH in the *Drosophila* Eye

To further characterize the morphological interaction between Myc and POSH, we used the GMR-GAL4 driver to restrict transgene expression to differentiating cells posterior to the morphogenetic furrow, thereby minimizing effects in earlier stages of eye development. Using this system, we first examined whether Myc overexpression alters *Posh* transcript abundance. RT-qPCR analysis of the eye imaginal discs expressing *UAS-Myc* under the control of *GMR-GAL4* revealed an approximately 2.4-fold increase in *Posh* mRNA levels relative to controls (Figure 4A). This finding shows that *Posh* transcript level increases upon Myc overexpression. We next examined adult eye morphology following overexpression of Myc and POSH individually or together (Figure 4B-E). While GMR-GAL4 eyes displayed a regular ommatidial arrangement (Figure 4B), Myc overexpression altered ommatidial organization (Figure 4C). In contrast, POSH overexpression produced a rough-eye phenotype with irregular ommatidial organization and disrupted surface architecture (Figure 4D). Co-expression of Myc and POSH resulted in a distinct and more pronounced disruption of the ommatidial pattern and eye surface morphology (Figure 4E). These findings demonstrate increased *Posh* transcript levels following Myc overexpression and support a genetic interaction affecting eye morphology.

**Figure 4.**
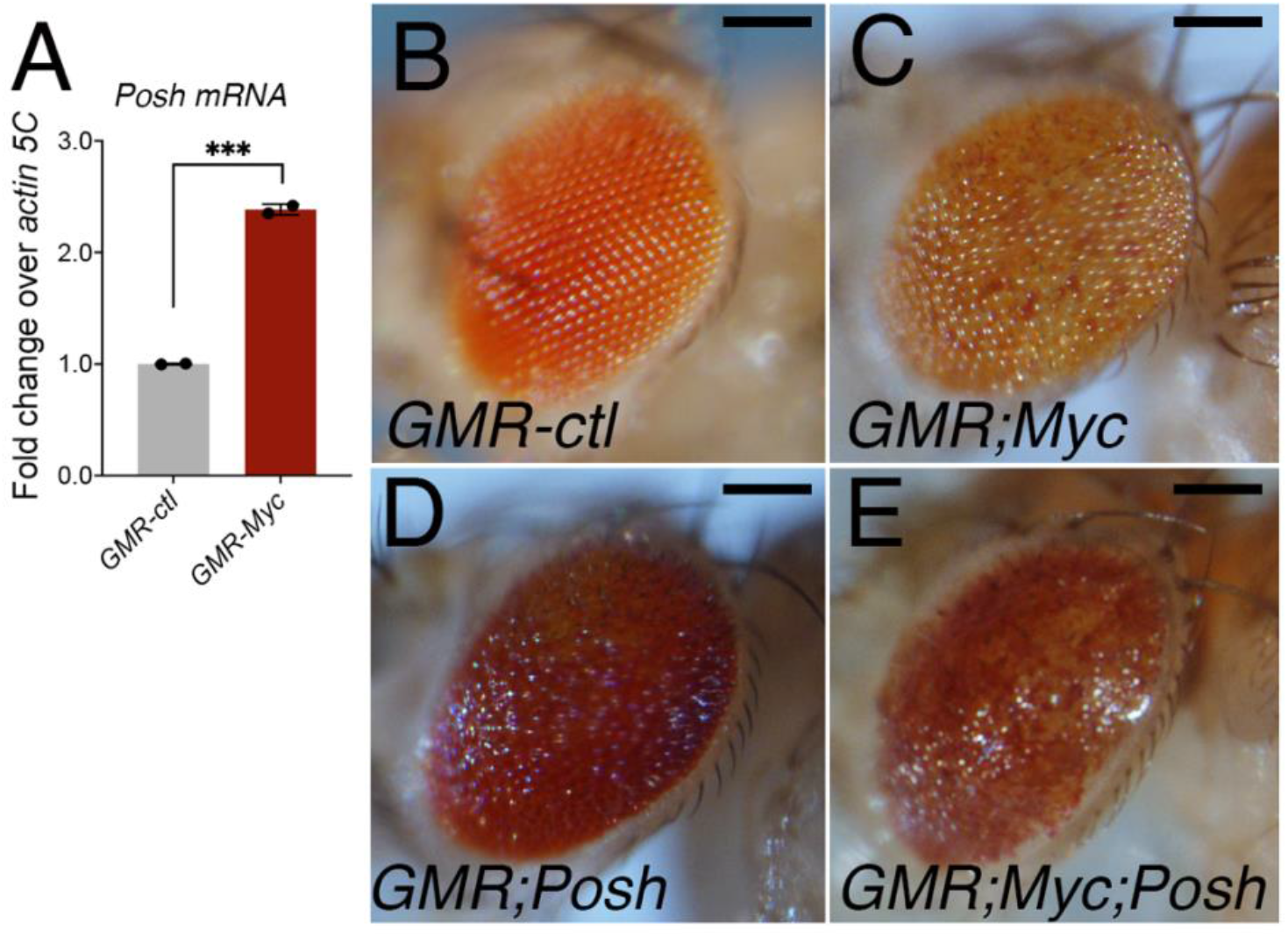
Myc overexpression increases *Posh* transcript levels and modifies adult eye morphology in combination with POSH. (A) RT-qPCR analysis of *Posh* mRNA in eye imaginal discs expressing *UAS-Myc* under the *GMR-GAL4* compared with *GMR-GAL4* controls. Transcript levels were normalized to *Actin5C* and expressed relative to the control. Data are mean ± SD from two independent experiments using approximately 30 eye-imaginal discs for genotype per experiment; ^***^*p* < 0.001. (B-E) Representative photographs of adult eyes expressing *GMR-GAL4* alone (B), Myc (C), POSH (D), and Myc and POSH together (E). Co-expression produced a pronounced alteration in the morphology of the eye surface architecture. Scale bars, 100 μm.

### 3.4 Myc and POSH Differentially Affect Ommatidial Size and Number

To further characterize the morphological interaction between Myc and POSH, we examined adult eyes overexpressing either Myc or POSH, individually or in combination, using the GMR-Gal4 promoter, and analyzed them by scanning electron microscopy (Figure 5A-D). Myc overexpression produced enlarged ommatidia, whereas POSH overexpression produced a rough-eye phenotype with irregular ommatidial organization. Co-expression of Myc and POSH caused a pronounced disruption of the regular ommatidial pattern. Quantification analysis revealed distinct effects on ommatidial size and number (Figure 5E-F). Myc overexpression increased mean ommatidial size area from 230 ± 5 μm^2^ in controls to 300 ± 5 μm^2^ (approximately 30%), without significantly affecting ommatidial number (690 ± 49 versus 652 ± 41). POSH overexpression had little effect on ommatidial area (230 ± 10 μm^2^) but reduced the total number to 579 ± 40 (approximately 16% below control). When Myc and POSH were co-expressed, ommatidial area remained elevated and did not differ significantly from that observed with Myc alone (Figure 5E). Conversely, Myc co-expression significantly attenuated the reduction in ommatidial number associated with POSH overexpression, bringing the count toward control values (Figure 5F). Thus, Myc primarily increased ommatidial size, whereas POSH predominantly reduced ommatidial number. Their co-expression caused substantial disorganization without a simple additive effect on either quantitative parameter.

**Figure 5.**
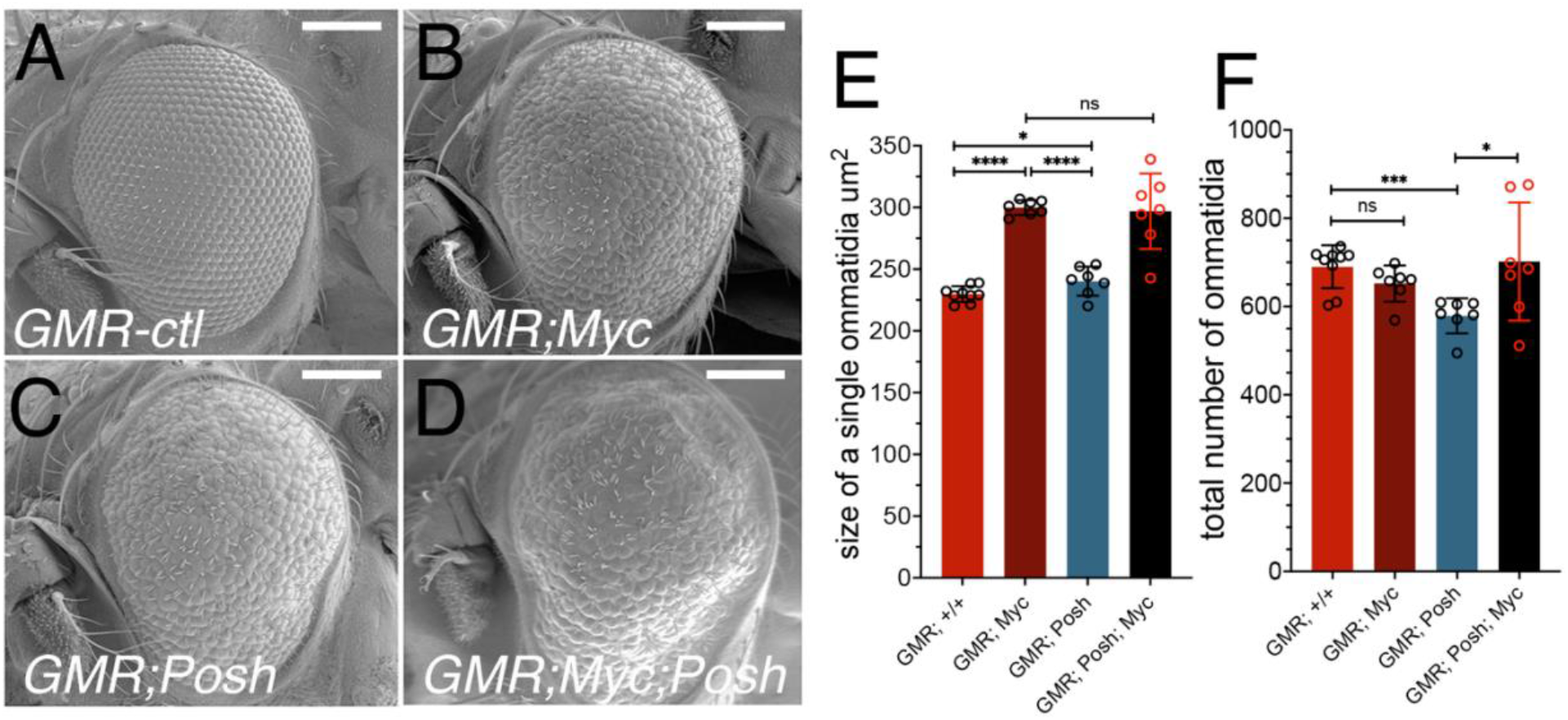
Myc and POSH differentially affect ommatidial size and number in the *Drosophila* eye. Representative scanning electron micrographs of adult eyes expressing *GMR-GAL4* alone (A), Myc (B), POSH (C), or Myc and POSH together (D). Quantification of mean ommatidial size (E) and total ommatidial number (F). Myc overexpression increased ommatidial size without significantly changing the number, whereas POSH overexpression reduced ommatidial number with little effect on ommatidial size. Co-expression of Myc and POSH maintained the Myc-associated increase in ommatidial area while attenuating the POSH-associated decrease in number. Data are mean ± SD; individual points represent independent eyes. Statistical significance is indicated in the graphs; ns, not significant; ^*^*p* < 0.05, ^***^*p* < 0.001, ^****^*p* < 0.0001. Scale bars, 100 μm.

To further examine the contribution of POSH to adult eye morphology and its genetic interaction with Myc, we analyzed the effects of reducing POSH using GMR-driven RNAi. POSH-RNAi alone did not produce obvious morphological changes relative to the luciferase RNAi control (Figure S1 A, B). Co-expression of POSH-RNAi attenuated the rough-eye phenotype induced by POSH overexpression (Figure S1 C, D). In contrast, reducing POSH in a Myc-overexpressing background did not visibly suppress the morphological increase in size of the ommatidia of Myc overexpression (Figure S2 A, B). These qualitative findings do not establish whether endogenous POSH is required for Myc-dependent growth, and further quantitative and functional analyses are needed to clarify its contribution.

## 4. Discussion

Here, we identify POSH as a genetic modifier of Myc function using the *Drosophila* eye. A genetic screen in a Myc-sensitized background identified the EP-1206 line, associated with increased *Posh* expression. This interaction was independently reproduced using a *UAS-POSH* transgene, supporting the conclusion that the EP-associated phenotype reflects altered POSH expression.

Progressively reducing Myc activity under the control of the *eyeless* compartment (Figure 3) resulted in smaller ommatidia and defects in head development, consistent with its established role in cellular growth [5,8]. Interestingly, the effects of POSH expression became progressively more pronounced as Myc activity decreased, resulting in increasingly severe eye phenotypes. This suggests that reduced Myc activity sensitizes developing eye tissue to POSH overexpression, potentially through an altered balance between growth-promoting and stress-responsive pathways. Further analysis using GMR-GAL4, which drives expression predominantly posterior to the morphogenetic furrow, where retinal differentiation is underway, revealed distinct effects of Myc and POSH on adult eye morphology and ommatidial organization (Figures 4 and 5). Myc overexpression increased ommatidial size without significantly affecting ommatidial number, whereas POSH overexpression primarily reduced ommatidial number and disrupted eye morphology. Both the reduction in ommatidial number and the rough-eye phenotype may partly reflect increased apoptosis, given the reported involvement of POSH in apoptotic signaling. Interestingly, co-expression of Myc and POSH led to a more severely disorganized eye phenotype, whereas Myc partially attenuated the reduction in ommatidial number caused by POSH overexpression. These apparently contrasting effects suggest that Myc and POSH influence distinct yet potentially interconnected processes that regulate cell growth, survival, and tissue organization. The increased *Posh* transcript levels observed following Myc overexpression (Figure 4A) further support a functional relationship between these regulators, although direct transcriptional regulation remains to be established. Using the GMR-GAL4 driver, we found that POSH reduction did not visibly modify the Myc-overexpression phenotype (Figures S1 and S2). Together with the effects of POSH overexpression in Myc-deficient backgrounds, these findings suggest that the genetic interaction may be particularly relevant under conditions of endogenous or reduced Myc activity, rather than Myc overexpression.

The genetic interaction between Myc and POSH may involve pathways regulating cellular growth, stress responses, and survival. POSH has been implicated in JNK-mediated apoptotic signaling [12,13,17,19,20,27], and Hippo/Yorkie regulation [21,28], both of which could contribute to the observed changes in ommatidial number and organization. Moreover, the increased *Posh* transcript levels following Myc overexpression, together with the presence of a putative E-box in its regulatory region [3], suggest a possible transcriptional relationship. However, whether these pathways contribute to the genetic interaction or *Posh* is directly regulated by Myc remains to be investigated.

## 5. Conclusions

In conclusion, our study identifies POSH as a novel genetic modifier of Myc function in the *Drosophila* eye and reveals its potential importance in coordinating cellular growth, stress responses, and tissue organization. Although the underlying mechanisms remain to be established, these findings provide a basis for further investigating the functional relationship between Myc and POSH during tissue development.

## Supplementary Materials

Figures S1 and S2 are provided with this article.

## Author Contributions

Conceptualization, P.B.; methodology, V.K., M.D., S.Z. and P.B.; validation, V.K., M.D., and S.Z.; formal analysis, V.K., M.D. and P.B.; investigation, V.K., M.D., and S.Z.; resources, P.B.; data curation, V.K., M.D., S.Z. and P.B.; writing original draft preparation, P.B.; writing, review, and editing, V.K., M.D., S.Z. and P.B.; project administration, P.B.; funding acquisition, P.B. All authors have read and agreed to the published version of the manuscript.

## Funding

This research was funded by the National Institute of Diabetes and Digestive and Kidney Diseases (NIDDK), National Institutes of Health (NIH), grant number SC1 DK085047, awarded to P.B. V.K. was supported by a fellowship from the CCNY/MSKCC Partnership for Cancer Research, Education (NCI), NIH-U54CA132378. The APC was waived by Genes.

## Institutional Review Board Statement

Not applicable. This study used Drosophila melanogaster, which is not subject to institutional animal ethical review requirements.

## Informed Consent Statement

Not applicable.

## Data Availability Statement

The data supporting the findings of this study are available within the article. Additional data are available from the corresponding author upon reasonable request.

## Acknowledgments

We thank the Bloomington Drosophila Stock Center (NIH P40OD018537) for providing Drosophila stocks. During the preparation of this manuscript, the authors used ChatGPT (OpenAI) for language editing and to improve the clarity of the text. The authors have reviewed and edited the output and take full responsibility for the content of this publication.

## Conflicts of Interest

The authors declare no conflicts of interest. The funders had no role in the design of the study; in the collection, analyses, or interpretation of data; in the writing of the manuscript; or in the decision to publish the results.

## Notes

### Competing Interest Statement

The authors have declared no competing interest.

## References

1. Gallant, P.,, Shiio, Y.,, Cheng, P.F.,, Parkhurst, S.M.,, Eisenman, R.N. Myc and Max homologs in Drosophila. Science 1996, 274, 1523–1527.

2. Bellosta, P.,, Gallant, P. Myc Function in Drosophila. Genes Cancer 2010, 1, 542–546, doi:10.1177/1947601910377490.

3. Hulf, T.,, Bellosta, P.,, Furrer, M.,, Steiger, D.,, Svensson, D.,, Barbour, A.,, Gallant, P. Whole-genome analysis reveals a strong positional bias of conserved dMyc-dependent E-boxes. Mol Cell Biol 2005, 25, 3401–3410.

4. Orian, A.,, Grewal, S.S.,, Knoepfler, P.S.,, Edgar, B.A.,, Parkhurst, S.M.,, Eisenman, R.N. Genomic binding and transcriptional regulation by the Drosophila Myc and Mnt transcription factors. Cold Spring Harb Symp Quant Biol 2005, 70, 299–307, doi:10.1101/sqb.2005.70.019.

5. Johnston, L.A.,, Prober, D.A.,, Edgar, B.A.,, Eisenman, R.N.,, Gallant, P. Drosophila myc regulates cellular growth during development. Cell 1999, 98, 779–790.

6. Destefanis, F.,, Manara, V.,, Bellosta, P. Myc as a Regulator of Ribosome Biogenesis and Cell Competition: A Link to Cancer. Int J Mol Sci 2020, 21, doi:10.3390/ijms21114037.

7. Grewal, S.S.,, Li, L.,, Orian, A.,, Eisenman, R.N.,, Edgar, B.A. Myc-dependent regulation of ribosomal RNA synthesis during Drosophila development. Nat Cell Biol 2005, 7, 295–302.

8. de la Cova, C.,, Abril, M.,, Bellosta, P.,, Gallant, P.,, Johnston, L.A. Drosophila myc regulates organ size by inducing cell competition. Cell 2004, 117, 107–116, doi:S0092867404002144 [pii].

9. Teleman, A.A.,, Hietakangas, V.,, Sayadian, A.C.,, Cohen, S.M. Nutritional control of protein biosynthetic capacity by insulin via Myc in Drosophila. Cell Metab 2008, 7, 21– 32, doi:S1550-4131(07)00365-8 [pii]10.1016/j.cmet.2007.11.010.

10. Parisi, F.,, Riccardo, S.,, Zola, S.,, Lora, C.,, Grifoni, D.,, Brown, L.M.,, Bellosta, P. dMyc expression in the fat body affects DILP2 release and increases the expression of the fat desaturase Desat1 resulting in organismal growth. Dev Biol 2013, 379, 64–75, doi:S0012-1606(13)00188-7 [pii]10.1016/j.ydbio.2013.04.008.

11. Parisi, F.,, Riccardo, S.,, Daniel, M.,, Saqcena, M.,, Kundu, N.,, Pession, A.,, Grifoni, D.,, Stocker, H.,, Tabak, E.,, Bellosta, P. Drosophila insulin and target of rapamycin (TOR) pathways regulate GSK3 beta activity to control Myc stability and determine Myc expression in vivo. BMC Biol 2011, 9, 65, doi:1741-7007-9-65 [pii]10.1186/1741-7007-9-65.

12. Montero, L.,, Muller, N.,, Gallant, P. Induction of apoptosis by Drosophila Myc. Genesis 2008, 46, 104–111.

13. Zhang, C.,, Casas-Tinto, S.,, Li, G.,, Lin, N.,, Chung, M.,, Moreno, E.,, Moberg, K.H.,, Zhou, L. An intergenic regulatory region mediates Drosophila Myc-induced apoptosis and blocks tissue hyperplasia. Oncogene 2015, 34, 2385–2397, doi:10.1038/onc.2014.160.

14. Nagy, P.,, Varga, A.,, Pircs, K.,, Hegedus, K.,, Juhasz, G. Myc-driven overgrowth requires unfolded protein response-mediated induction of autophagy and antioxidant responses in Drosophila melanogaster. PLoS Genet 2013, 9, e1003664, doi:10.1371/journal.pgen.1003664PGENETICS-D-13-00114 [pii].

15. Paiardi, C.,, Mirzoyan, Z.,, Zola, S.,, Parisi, F.,, Vingiani, A.,, Pasini, M.E.,, Bellosta, P. The Stearoyl-CoA Desaturase-1 (Desat1) in Drosophila cooperated with Myc to Induce Autophagy and Growth, a Potential New Link to Tumor Survival. Genes (Basel) 2017, 8, doi:10.3390/genes8050131.

16. Moreno, E.,, Basler, K. dMyc transforms cells into super-competitors. Cell 2004, 117, 117–129.

17. Tapon, N.,, Nagata, K.,, Lamarche, N.,, Hall, A. A new rac target POSH is an SH3- containing scaffold protein involved in the JNK and NF-kappaB signalling pathways. EMBO J 1998, 17, 1395–1404, doi:10.1093/emboj/17.5.1395.

18. Wilhelm, M.,, Kukekov, N.V.,, Xu, Z.,, Greene, L.A. Identification of POSH2, a novel homologue of the c-Jun N-terminal kinase scaffold protein POSH. Dev Neurosci 2007, 29, 355–362, doi:000105476 [pii]10.1159/000105476.

19. Lennox, A.L.,, Stronach, B. POSH misexpression induces caspase-dependent cell death in Drosophila. Dev Dyn 2010, 239, 651–664, doi:10.1002/dvdy.22186.

20. Tsuda, M.,, Seong, K.H.,, Aigaki, T. POSH, a scaffold protein for JNK signaling, binds to ALG-2 and ALIX in Drosophila. FEBS Lett 2006, 580, 3296–3300, doi:10.1016/j.febslet.2006.05.005.

21. Ma, X.,, Guo, X.,, Richardson, H.E.,, Xu, T.,, Xue, L. POSH regulates Hippo signaling through ubiquitin-mediated expanded degradation. Proc Natl Acad Sci U S A 2018, 115, 2150–2155, doi:10.1073/pnas.1715165115.

22. Bellosta, P.,, Hulf, T.,, Balla Diop, S.,, Usseglio, F.,, Pradel, J.,, Aragnol, D.,, Gallant, P. Myc interacts genetically with Tip48/Reptin and Tip49/Pontin to control growth and proliferation during Drosophila development. Proc Natl Acad Sci U S A 2005, 102, 11799–11804, doi:0408945102 [pii]10.1073/pnas.0408945102.

23. Rorth, P. A modular misexpression screen in Drosophila detecting tissue-specific phenotypes. Proc Natl Acad Sci U S A 1996, 93, 12418–12422, doi:10.1073/pnas.93.22.12418.

24. Rorth, P.,, Szabo, K.,, Bailey, A.,, Laverty, T.,, Rehm, J.,, Rubin, G.M.,, Weigmann, K.,, Milan, M.,, Benes, V.,, Ansorge, W.,, et al. Systematic gain-of-function genetics in Drosophila. Development 1998, 125, 1049–1057, doi:10.1242/dev.125.6.1049.

25. Karkkainen, S.,, van der Linden, M.,, Renkema, G.H. POSH2 is a RING finger E3 ligase with Rac1 binding activity through a partial CRIB domain. FEBS Lett 2010, 584, 3867– 3872, doi:10.1016/j.febslet.2010.07.060.

26. Seong, K.H.,, Matsuo, T.,, Fuyama, Y.,, Aigaki, T. Neural-specific overexpression of drosophila plenty of SH3s (DPOSH) extends the longevity of adult flies. Biogerontology 2001, 2, 271–281, doi:10.1023/a:1013249326285.

27. Xu, Z.,, Kukekov, N.V.,, Greene, L.A. POSH acts as a scaffold for a multiprotein complex that mediates JNK activation in apoptosis. EMBO J 2003, 22, 252–261, doi:10.1093/emboj/cdg021.

28. Xiao, W.,, Wang, J.,, Ou, C.,, Zhang, Y.,, Ma, L.,, Weng, W.,, Pan, Q.,, Sun, F. Mutual interaction between YAP and c-Myc is critical for carcinogenesis in liver cancer. Biochem Biophys Res Commun 2013, 439, 167–172, doi:S0006-291X(13)01418-6 [pii]10.1016/j.bbrc.2013.08.071.

