## Supplementary for "POSH (Plenty of SH3): a novel genetic modifier of Myc function in Drosophila"

**Supplementary Figures:**

**Supplementary Figure S1.**

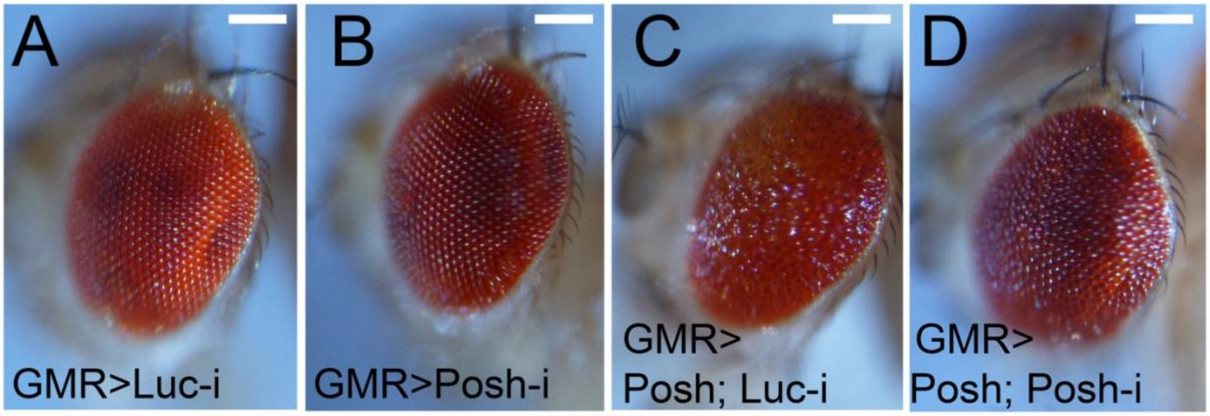


**Effects of reducing Posh on adult *Drosophila* eye morphology and in Posh-overexpression.** Representative light microscopy images of adult *Drosophila* eyes expressing the indicated transgenes under the control of the GMR-Gal4 driver. **(A)** Control eye expressing luciferase RNAi (GMR>Luc-i). **(B)** Posh knockdown (GMR>Posh-i) does not produce obvious alterations in eye size or ommatidial organization compared with the control. **(C)** Posh overexpression (GMR>Posh; Luc-i) induces alterations in eye morphology and ommatidial organization. **(D)** Co-expression of Posh RNAi (GMR>Posh; Posh-i) attenuates the morphological defects associated with Posh overexpression. Scale bars: 100 µm.

**Supplementary Figure S2.**

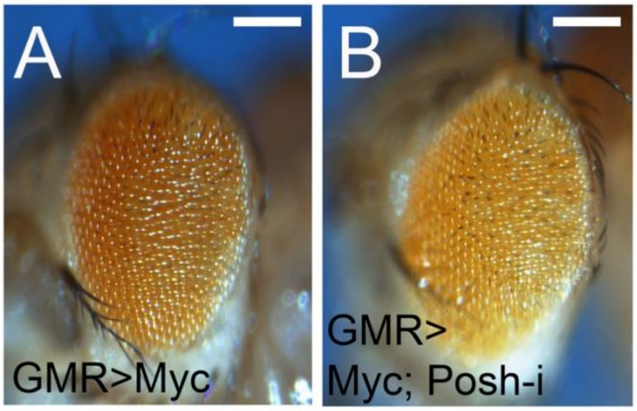


**Effect of Posh knockdown on Myc-induced eye phenotypes.** Representative light microscopy images of adult *Drosophila* eyes expressing the indicated transgenes under the control of the GMR-Gal4 driver. **(A)** Myc overexpression (GMR>Myc). **(B)** Co-expression of Myc and Posh RNAi (GMR>Myc; Posh-i) does not produce an obvious modification of the Myc-induced eye phenotype. Indeed,Posh knockdown does not visibly suppress the Myc-overexpression phenotype, although this qualitative observation does not exclude more subtle effects. The GMR-Gal4 control eye is shown in Supplementary Figure S1A (GMR>Luc-i). Scale bars: 100 µm.
